# Efficacy and safety of a monodisperse solid-in-oil-in-water emulsion in transcatheter arterial chemoembolization in a rabbit VX2 tumor model

**DOI:** 10.1101/759316

**Authors:** Daisuke Yasui, Aya Yamane, Hiroshi Itoh, Masayuki Kobayashi, Shin-ichiro Kumita

## Abstract

Transcatheter arterial chemoembolization (TACE) is a standard treatment for unresectable hepatocellular carcinoma; however, it does not always result in tumor control. Nevertheless, treatment outcome can be improved with monodisperse emulsions of anticancer agents. In this study, the efficacy and safety of a monodisperse miriplatin-Lipiodol emulsion were evaluated in Japanese white rabbits. VX2 tumor was implanted into the left liver lobe of each rabbit. The animals were divided into control and experimental groups (of five animals each) and respectively administered a conventional miriplatin suspension or the emulsion via the left hepatic artery. Computed tomography (CT) was performed before, immediately after, and two days following TACE. All rabbits were sacrificed two days after the procedure. Each tumor was removed and cut in half for assessment of iodine concentration in one half by mass spectroscopy and evaluation of Lipiodol accumulation and adverse events in the other half. Mean Hounsfield unit (HU) values were measured using plain CT images taken before and after TACE. Iodine concentration was higher in the experimental group [1100 (750–1500) ppm] than in the control group [840 (660–1800) ppm]. Additionally, the HU value for the experimental group was higher than that for the control group immediately after [199.6 (134.0– 301.7) vs. 165.3 (131.4–280.5)] and two days after [114.2 (56.1–229.8) vs. 58.3 (42.9–132.5)] TACE. Cholecystitis was observed in one rabbit in the control group. Ischemic bile duct injury was not observed in any group. The results show that Lipiodol accumulation and retention in VX2 tumor may be improved by using a monodisperse emulsion. Moreover, no significant adverse events are associated with the use of the emulsion.

## Introduction

Transcatheter arterial chemoembolization (TACE) is a standard of care for unresectable hepatocellular carcinoma (HCC), according to the Barcelona Clinic Liver Cancer (BCLC) treatment protocol.[1] The procedure consists of two steps. First, a mixture of anticancer agents (doxorubicin, epirubicin, cisplatin, and miriplatin) and iodized oil (Lipiodol; Guerbet, Villepinte, France) is injected through microcatheters placed in a feeding artery. Gelatin slurry is subsequently injected to induce ischemia. The efficacy of TACE has been validated in randomized control trials.[2] There have been several attempts to further improve local tumor control by using microspheres and performing balloon-occluded TACE; however, complete response rates remain in the range of 26.9–57.1 %.[3–6]

Two types of carriers are now available for transarterial delivery of anticancer agents: Lipiodol and microspheres. Well to moderately-differentiated HCC is usually fed by abnormal arteries that do not accompany bile ducts (unpaired arteries) and are drained by terminal portal venules or adjacent hepatic sinusoids,[7] which can be potential routes for metastasis. Lipiodol is a liquid that can be distributed to the drainage area, while microspheres cannot. Therefore, Lipiodol is considered a better embolization agent compared to microspheres for obtaining local tumor control.

Unsatisfactory outcomes of TACE using Lipiodol may be due to uneven sizes of the anticancer agent-Lipiodol emulsion particles. Consequently, using a monodisperse anticancer agent-Lipiodol emulsion may improve local tumor control. Emulsion has been prepared by manually mixing two agents using a three-way stopcock without adding a surfactant. The resultant polydisperse emulsion is unstable and will typically get stuck in the proximal feeding arteries, thus compromising the delivery of an anticancer agent to the tumor. A solution to this problem is to use a monodisperse emulsion, which can be prepared by membrane emulsification using the following procedure. A mixture of the anticancer agent and Lipiodol (water-in-oil emulsion, w/o emulsion) is pushed under constant pressure through an SPG membrane (from SPG Technology Co., Ltd.), which has multiple even-sized pores. A homogenous emulsion, specifically a water-in-oil-in-water (w/o/w) emulsion, can then be obtained. Clinical application of this type of dispersion has been already attempted using epirubicin w/o/w emulsion and promising results have been obtained.[8] However, the process for preparing a w/o/w emulsion is complex. Moreover, emulsions that are prepared using lipophilic anticancer agents are more stable since their affinity for Lipiodol is higher than that of hydrophilic agents.

Miriplatin is a lipophilic platinum derivative.[9] It is prepared as a freeze-dried powder, which can be readily dispersed in Lipiodol as a suspension. A solid-in-oil-in-water (s/o/w) emulsion of miriplatin can be obtained by pushing a miriplatin-Lipiodol suspension through an SPG membrane. The aim of the present study was to evaluate the efficacy and safety of miriplatin s/o/w emulsion and compare them to those of a conventional suspension in Japanese white rabbits.

## Materials and Methods

### Preparation of conventional miriplatin suspension and miriplatin s/o/w emulsion

Miriplatin (Sumitomo Dainippon Pharma Co., Ltd., Osaka, Japan) suspension was prepared using 60 mg miriplatin powder and 3 mL Lipiodol. In order to prepare the emulsion, the suspension was pushed through a 20-μm hydrophilic SPG membrane (SPG Technology Co., Ltd., Miyazaki, Japan) into an outer aqueous phase using a syringe pump (https://dx.doi.org/10.17504/protocols.io.6edhba6). PEG-60 hydrogenated castor oil (HCO 60; Nikko Chemicals Co., Ltd., Tokyo, Japan) was used as a surfactant in the formulation. HCO 60 (0.8 %) was added to NaCl solution (0.45 %w/v) to obtain a mixture that was used as the outer aqueous phase. Droplet size was measured by microscopy at 100× magnification. Kernel density estimation was performed to estimate the distribution of droplet diameter.

### Animal experiment

The study was carried out in strict accordance with the relevant guidelines and acts in Japan, including the Act on Welfare and Management of Animals. The protocol was approved by the Animal Experiment Ethics Committee of Nippon Medical School (Tokyo, Japan; protocol number: 30-014). All efforts were made to minimize suffering. VX2-tumor-bearing Japanese white rabbits (14 weeks old, clean; Japan SLC, Inc., Hamamatsu, Japan) were used for the study. The tumor was implanted in the left liver lobe of each rabbit. TACE was performed 14 days after tumor implantation. Five rabbits each were put in control and experimental groups and administered the conventional suspension and s/o/w emulsion, respectively.

Dynamic contrast-enhanced computed tomography (CT) scan (Aquilion PRIME; Canon Medical Systems, Tochigi, Japan) was performed prior to TACE. The contrast medium (iopamidol, 300 mg I/mL; Fuji Pharma Co., Ltd., Tokyo, Japan) was intravenously injected at 2 mL/kg at a rate of 0.2 mL/s into the animals. Quadriphasic scan was performed at 20, 40, 63, and 118 s after the injection. The scanning parameters used were as follows: collimation thickness, 80 × 0.5 mm; rotation speed, 0.35 s; helical pitch, 0.738; tube voltage, 80 kVp; and tube current, automatic exposure control.

TACE was performed under general anesthesia, which was induced using a subcutaneous injection of medetomidine hydrochloride (0.1 mg/kg) and ketamine hydrochloride (25 mg/kg) before moving to angio suite. Anesthesia was maintained by inhalation of 2–2.5 % isoflurane in angio suite. Each rabbit was monitored during the procedure using an electrocardiograph and a pulse oximeter. The common femoral artery was surgically exposed, into which a 4-F introducer sheath (Medikit Co., Ltd., Tokyo, Japan) was inserted. The celiac artery was cannulated with a 4-F C2 diagnostic catheter (Medikit Co., Ltd.). Next, a 2-F microcatheter (Gold Crest Neo; HI-LEX, Takarazuka, Japan) was coaxially advanced into the left hepatic artery, after which 0.1 mL of either the suspension or emulsion was injected under fluoroscopic guidance. A plain CT scan was taken immediately after the procedure.

All the rabbits were euthanized two days after TACE by injecting pentobarbital sodium (100mg/kg) intravenously. Dynamic contrast-enhanced CT scanning was performed prior to sacrifice by following the same protocol as before TACE. The tumor was extracted and cut in half: one half was used for pathological evaluation, whereas the other half was evaluated for its iodine content by mass spectroscopy.

### Evaluation of the efficacy of the emulsion

Accumulation of iodized oil in the tumors was assessed by mass spectroscopy (for iodine concentration) as well as pathological and radiological analyses.

#### Mass spectroscopy

Iodine concentration in the specimens was evaluated by inductively-coupled plasma mass spectroscopy (Japan Testing Laboratories, Inc., Ogaki, Japan). Median iodine concentration was then compared between the two animal groups.

#### Pathological evaluation

Oil red O staining, which is a lipid staining test, was performed to quantitatively evaluate the accumulation of iodized oil in the tumors. Each specimen was prepared using the maximum cross-section of the tumor. Digitization was done by whole slide imaging (NanoZoomer-XR; Hamamatsu Photonics K.K., Yokohama, Japan). Each scanned image was compressed to obtain a JPEG image with a resolution of 1920 × 984 pixels. Lipiodol accumulation was evaluated using the HSV model, which defines color space by three parameters: hue, saturation, and value. Lesions without lipid were bluish in color. An area with a ‘hue’ value of more than 55 and less than 65 was digitally subtracted from the digitized specimen by setting ‘value’ to be zero. The remaining area was considered to be the region of Lipiodol accumulation. The extent of Lipiodol accumulation was evaluated using ‘saturation’. Source codes were written in Python (Anaconda 4.6.14; Anaconda, Inc., Austin, TX, USA) using Jupyter Notebook, whereas cv2 module was used for image processing as described above.

#### Radiological evaluation

A slice of specimen was selected to include the maximum cross-section of the tumor for the evaluation. A region of interest (ROI) was defined to cover the target tumor in the CT images taken before, immediately after (POD 0), and two days after (POD 2) TACE. Mean Hounsfield unit (HU) values were used for the evaluation. Washout rate was defined as the difference in HU values between POD 0 and POD 2, divided by the HU value for POD 0.

### Evaluation of adverse events

Cholecystitis, bile duct injury, and biloma formation were considered as potential adverse events. The extent of Lipiodol accumulation in the gallbladder was evaluated using CT images, whereas inflammation of the gallbladder was evaluated using pathological specimens. Bile duct injury and biloma formation were pathologically evaluated.

## Results

A monodisperse s/o/w emulsion containing miriplatin was successfully formulated with a mean diameter of 62.0 ± 6.42 μm (Fig 1). Kernel density estimation showed that the mode droplet size was 60.0 μm. All experimental procedures were successfully performed.

**Fig 1.**
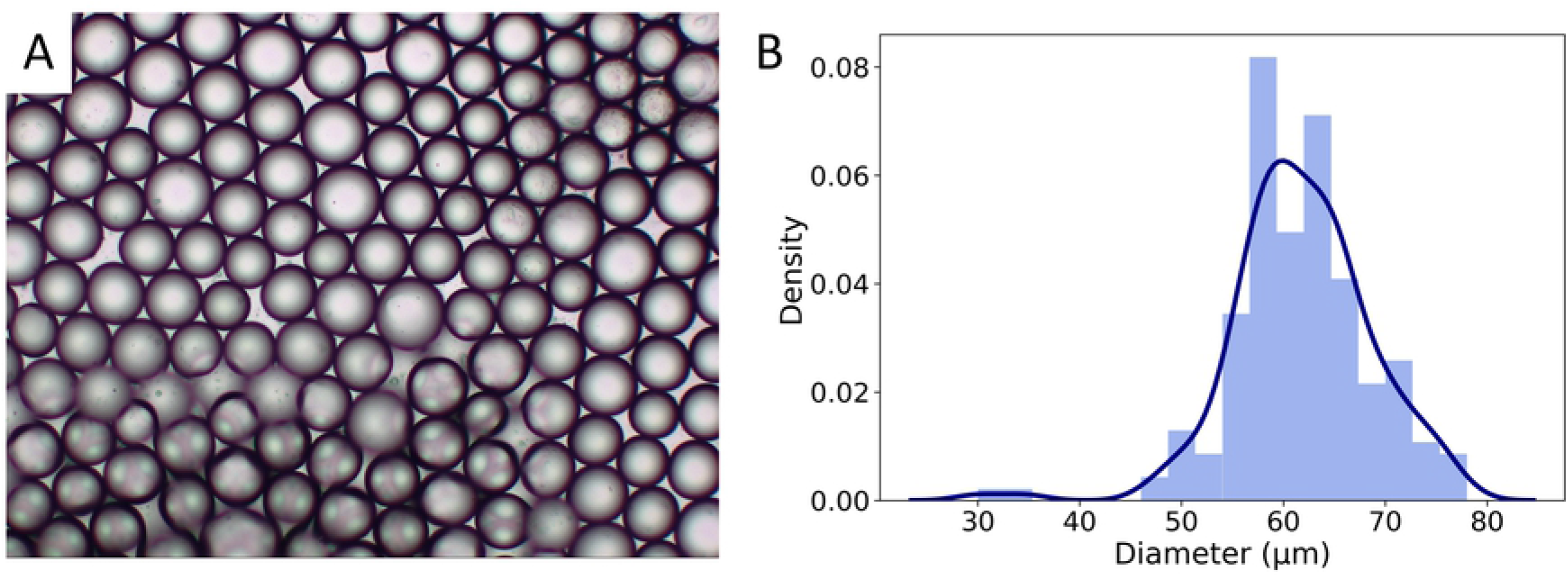
The s/o/w miriplatin emulsion. (A) Image of s/o/w miriplatin emulsion (as obtained from the microscopic analysis) (100× magnification). (B) Results of kernel density estimation. The data was calculated based on droplet diameters obtained from the microscopic analysis.

The mean body weights of the rabbits were 2.6 ± 0.1 and 2.7 ± 0.1 kg in the control and experimental groups, respectively. Tumor diameter was 14.3 ± 5.6 mm in the control group and 15.4 ± 3.0 mm in the experimental group. All tumors were confirmed to be in the left lobe by CT scan prior to performing TACE.

Median iodine concentration was higher in the experimental group (1100 ppm, range: 750– 1500 ppm) than in the control group (840 ppm, range: 660–1800 ppm (Fig 2). The median saturation was also higher in the experimental group (4.4, range: 2.8–20.3) than in the control group (3.8, range: 2.1–8.1) (Fig 3). Mean HU value was higher in the experimental group than in the control group immediately after TACE [199.6 (134.0–301.7) vs. 165.3 (131.4–280.5)] as well as two days after TACE [114.2 (56.1–229.8) vs. 58.3 (42.9–132.5)] (Fig 4). Washout rate was higher in the control group [55.6 (51.1–75.4)] than in the experimental group [49.9 (23.8–58.1)] (Fig 4). Lipiodol accumulation varied greatly even among animals in the same group. Extensive Lipiodol deposition was observed in all but one rabbit in the experimental group (Figs 5 and 6). Poor Lipiodol deposition was possibly due to a massive arterioportal shunt around the tumors. Lipiodol tended to deposit in normal liver parenchyma rather than in the tumors, as observed in the pathological specimens and CT images (Fig 6). A low-density area was observed at the interface between the tumor and the normal liver parenchyma, which possibly indicates liquefying necrosis, in all the rabbits in the experimental group and in three rabbits in the control group (Figs 5 and 6).

**Fig 2.**
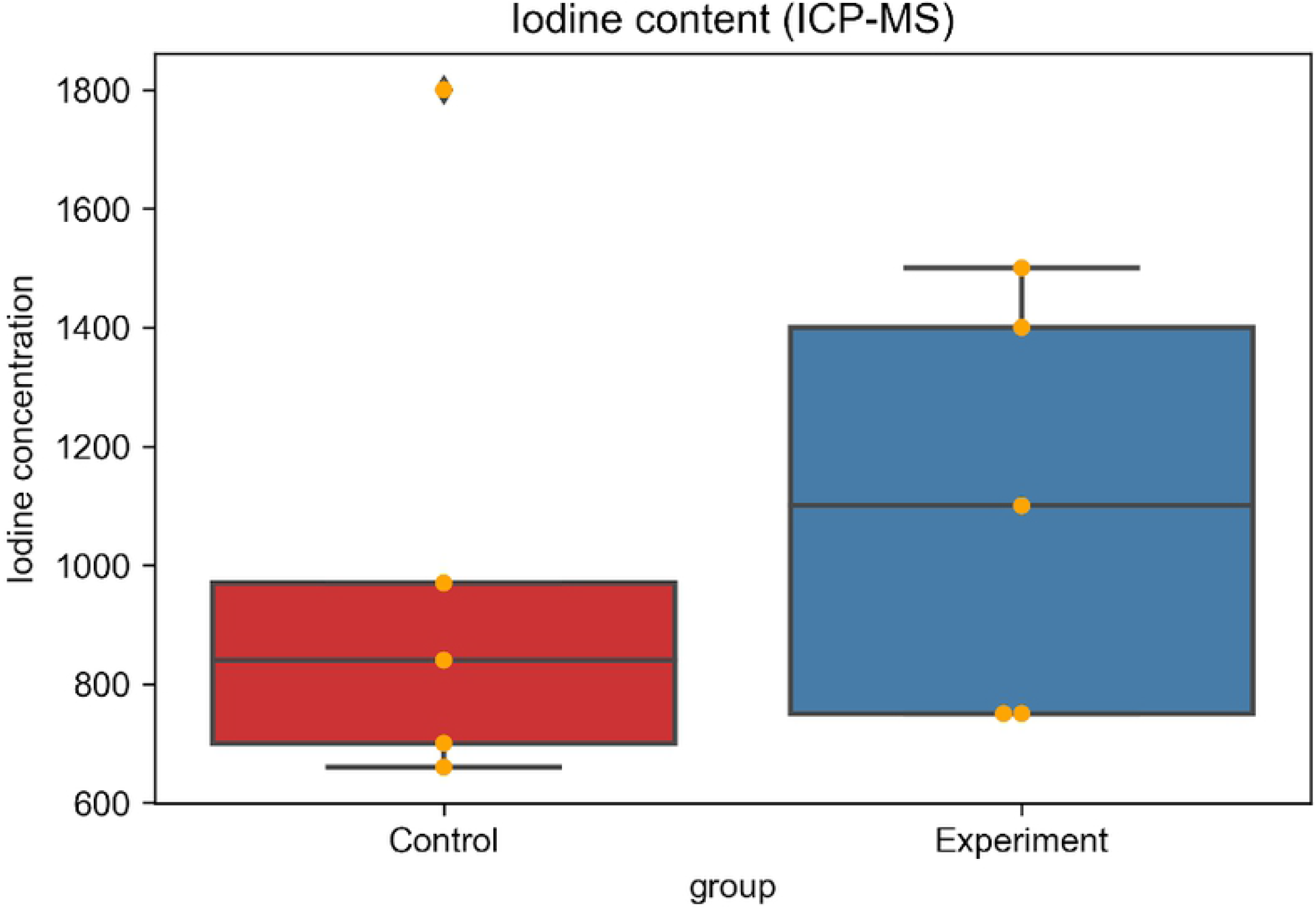
Iodine concentration in the tumors. The concentration of iodine (ppm) in the tumors was evaluated by inductively-coupled plasma mass spectroscopy.

**Fig 3.**
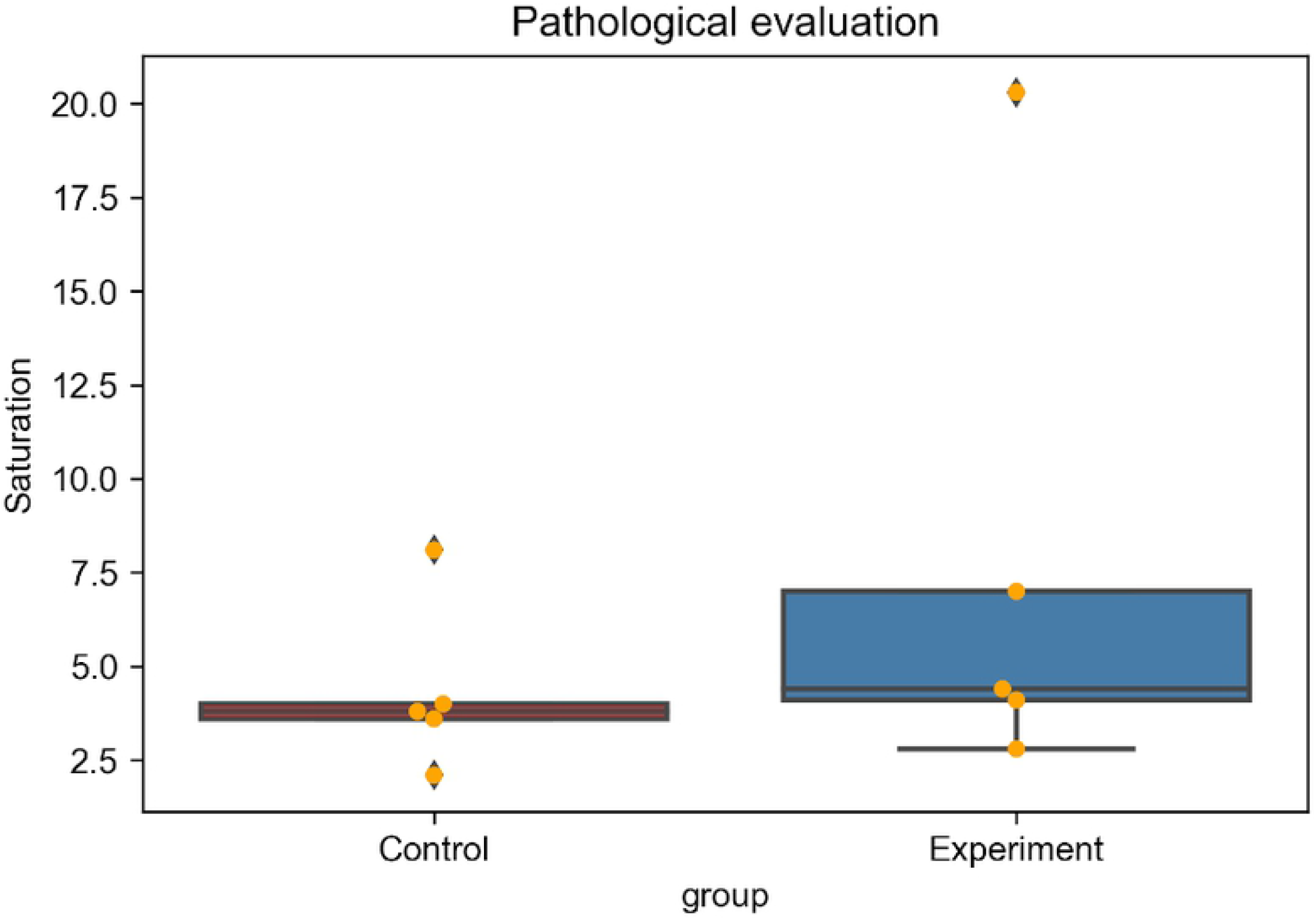
Quantitative pathological evaluation of Lipiodol accumulation in the tumors. The HSV model was used in the assessment. ‘Saturation’ values for areas with Lipiodol accumulation were evaluated using digitized specimens stained with Oil red O.

**Fig 4.**
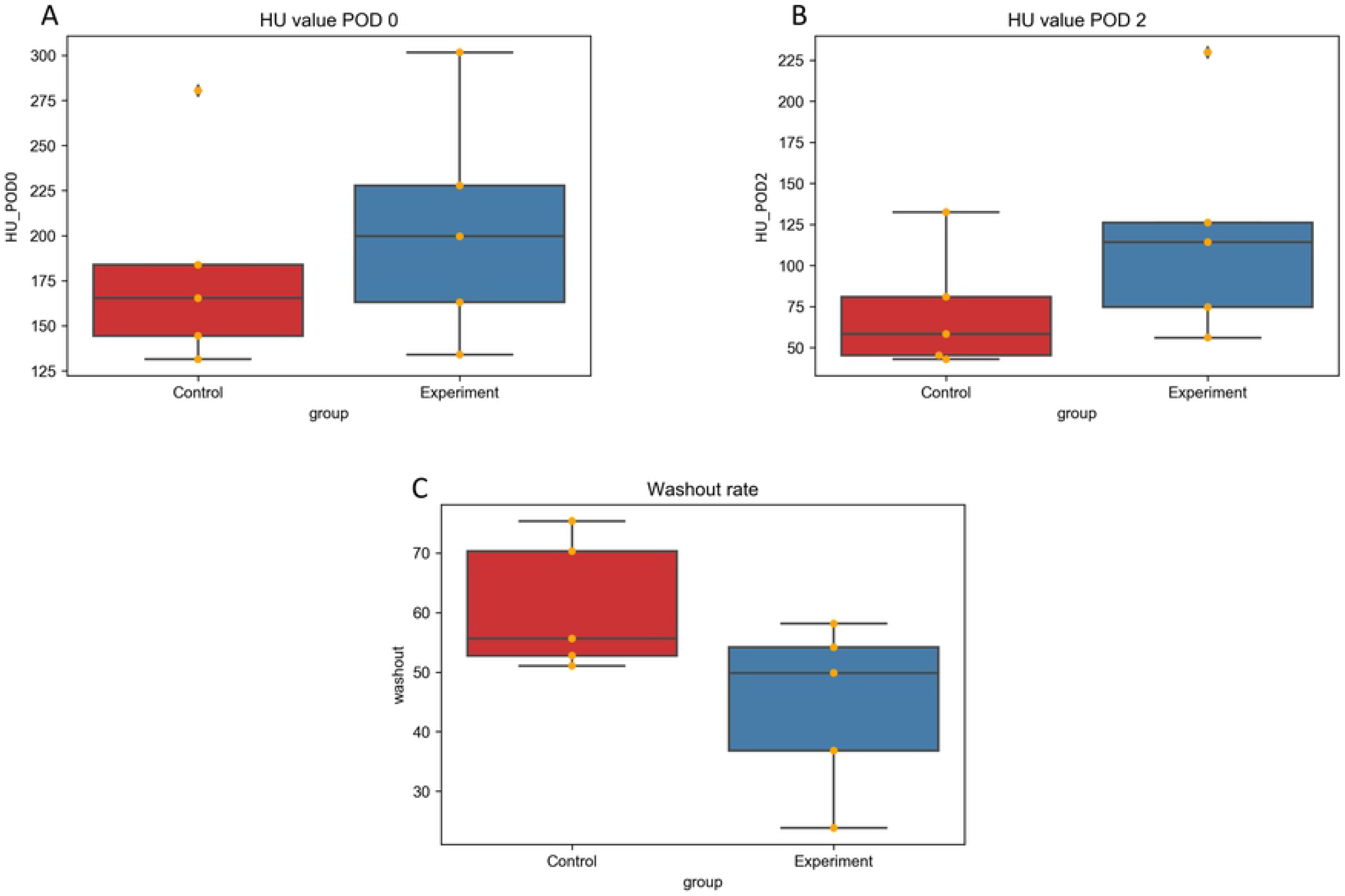
Radiological evaluation of Lipiodol accumulation in the tumors. Mean HU value (A) immediately after and (B) two days after TACE. (C) Lipiodol washout rate.

**Fig 5.**
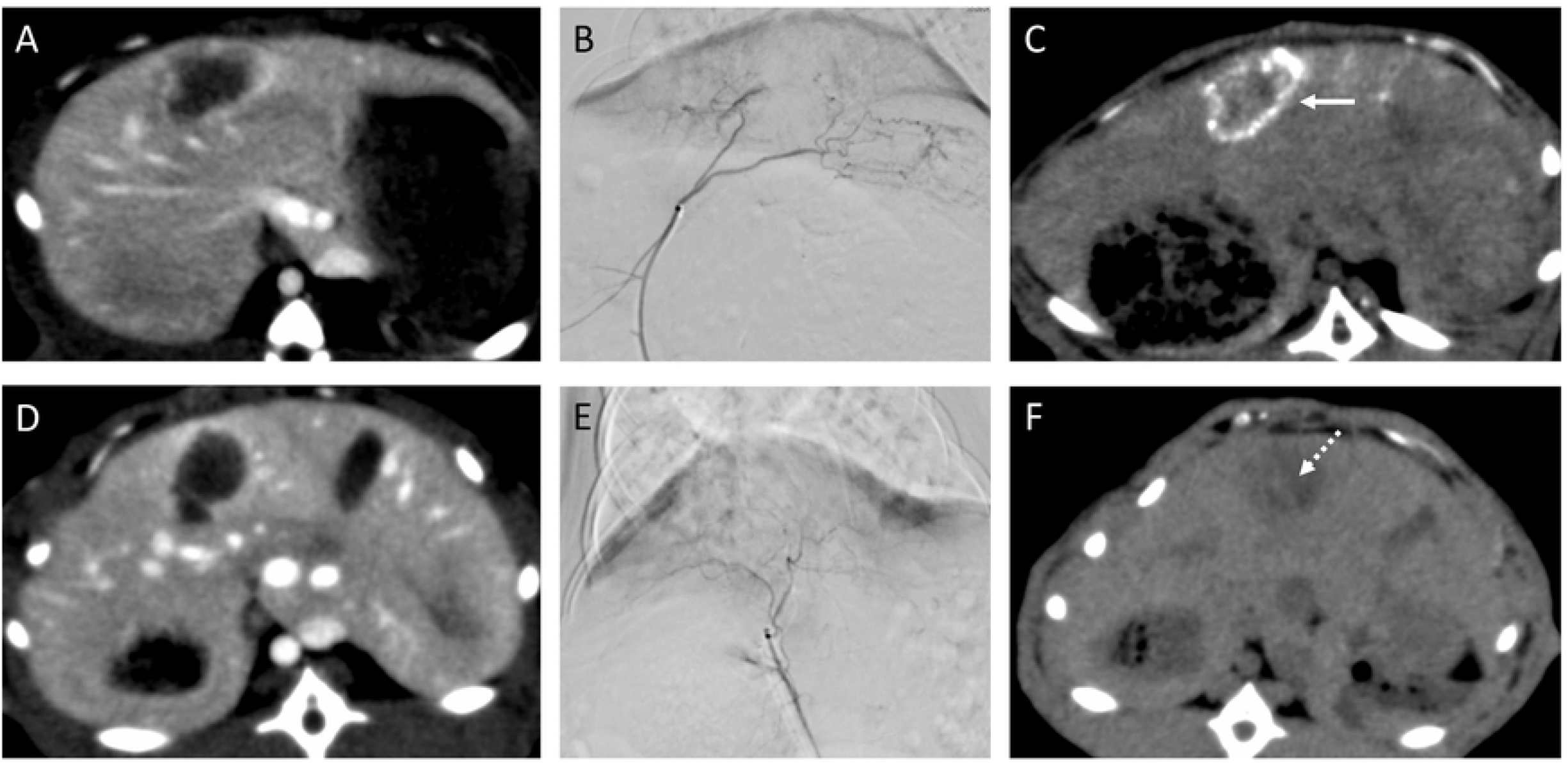
Representative images showing Lipiodol deposition. (A–C) Images for a rabbit in the experimental group. The s/o/w miriplatin emulsion was injected via the left hepatic artery. (C) Dense Lipiodol accumulation observed two days after TACE (white arrow). (D–F) Images for a rabbit in the control group. Miriplatin suspension was injected via the left hepatic artery. (F) Slight accumulation of Lipiodol observed two days after TACE (white dashed arrow).

**Fig 6.**
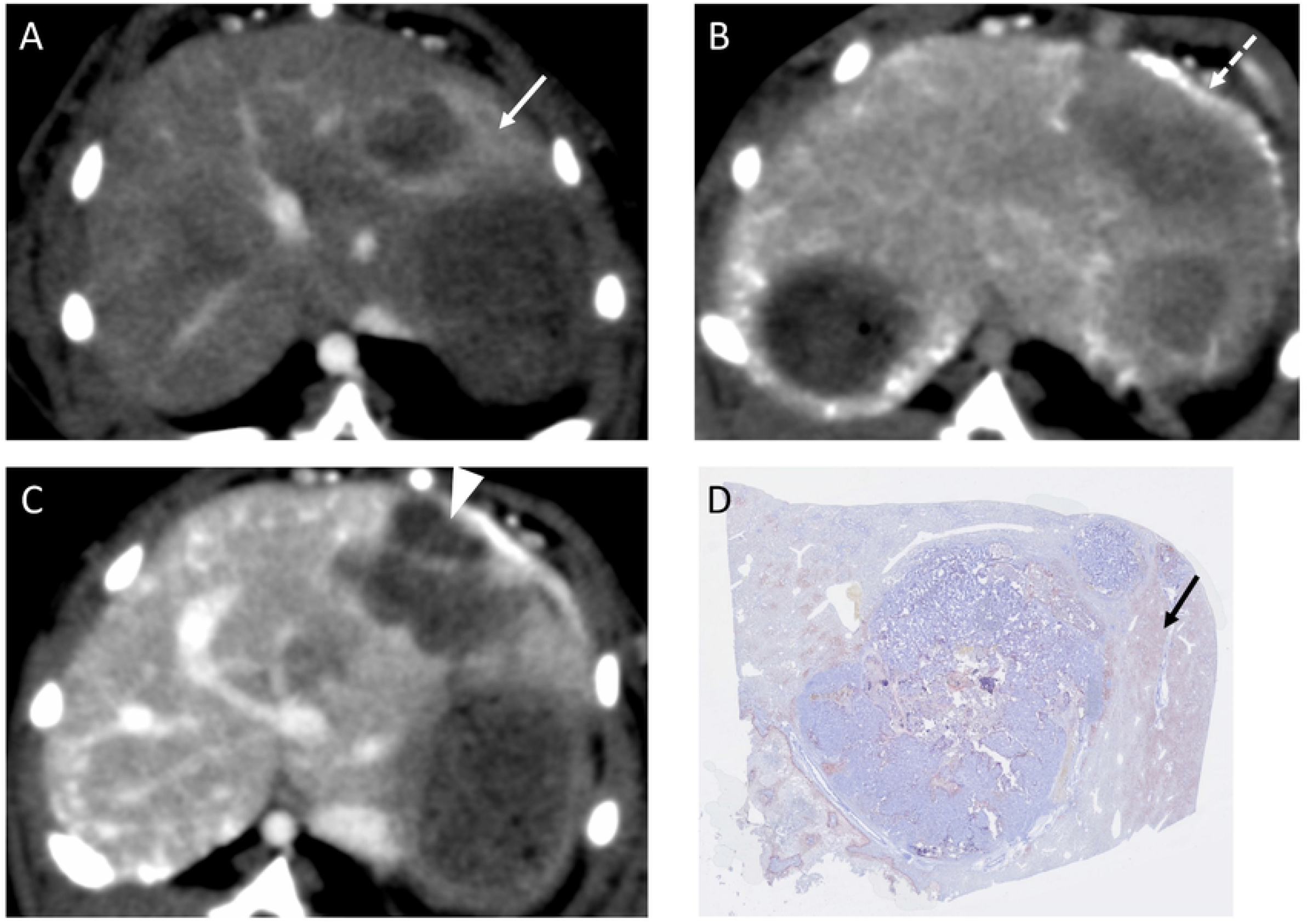
Representative images showing poor Lipiodol accumulation in a rabbit in the experimental group. (A) A massive arterioportal shunt around the tumor (white arrow). (B) Lipiodol accumulation higher in the liver parenchyma than in the tumor immediately after TACE (white dashed arrow). (C) Necrosis in liver parenchyma surrounding the tumor two days after TACE (white arrowhead). (D) Lipiodol accumulation in normal liver parenchyma in specimen stained with Oil red O (black arrow).

In the experimental group, Lipiodol accumulation in the gallbladder was observed in four rabbits immediately after TACE; however, washout was observed in all rabbits and cholecystitis was not observed. Conversely, in the control group, Lipiodol accumulation was observed in three rabbits after TACE and washout was not observed in one animal, in which cholecystitis ensued (Fig 7). Bile duct injury and biloma were not observed in either group.

**Fig 7.**
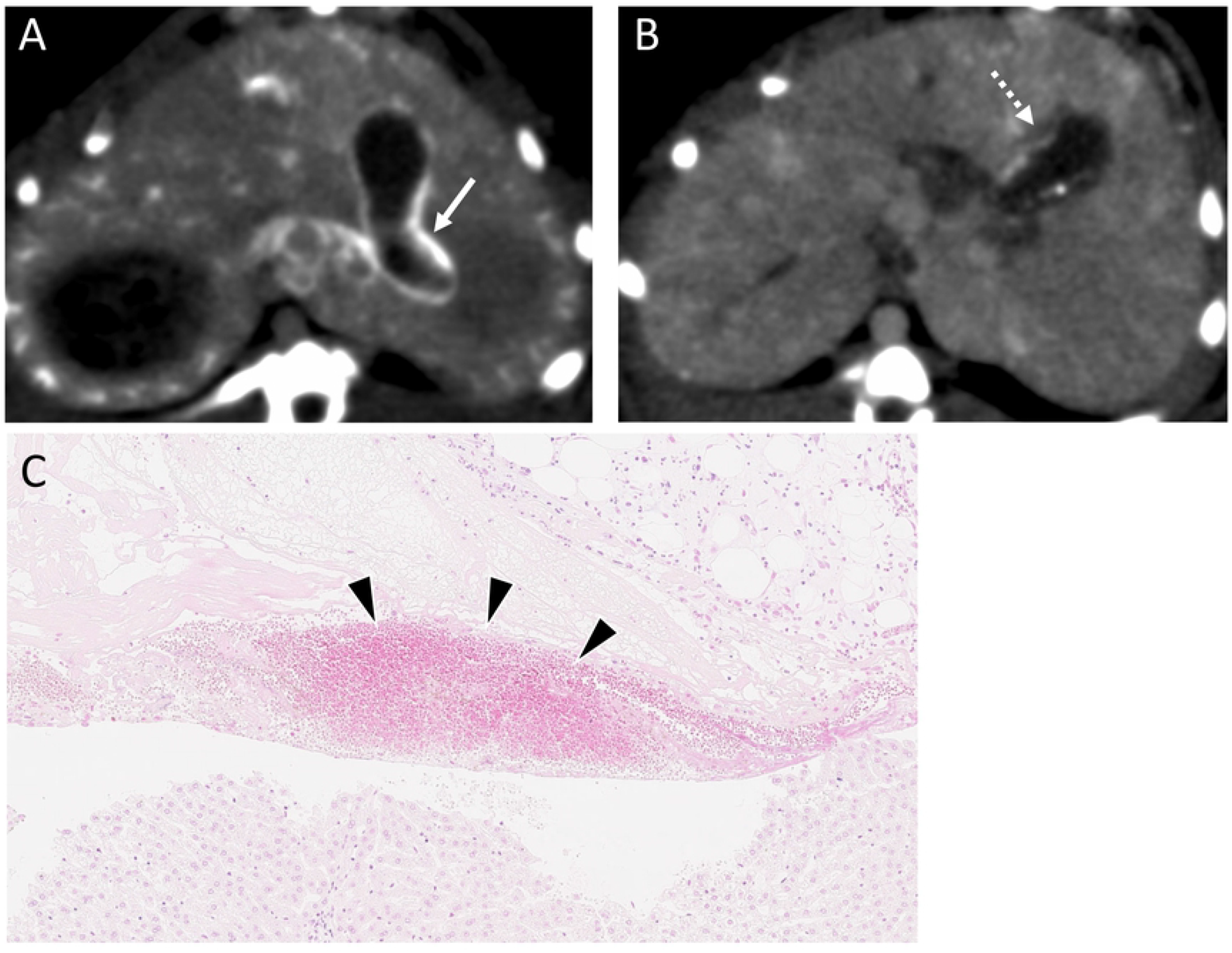
A case of cholecystitis following TACE. (A) Diffuse Lipiodol accumulation in the gallbladder wall, as observed in a CT scan, immediately after TACE (white arrow). (B) Lipiodol retention in the gallbladder wall, as well as thickening of the wall, two days after TACE (white dashed arrow). (C) Infiltration of inflammatory cells into the gallbladder wall, which is indicative of cholecystitis (black arrowheads).

## Discussion

Improved accumulation of Lipiodol in the VX2 tumors was observed using the novel s/o/w miriplatin emulsion, compared to the conventional miriplatin suspension. It is considered that Lipiodol distribution to the tumor was improved because a monodisperse emulsion was used. This was verified by performing mass spectroscopy, radiological, and pathological analyses. Miriplatin is a lipophilic platinum derivative, which was specially designed to have an increased affinity for Lipiodol. Therefore, it can be well dispersed in Lipiodol to form an aqueous suspension and gradually release active platinum into the aqueous phase.[10] Miriplatin has several advantages over other anticancer drugs such as doxorubicin, epirubicin, cisplatin, and mitomycin C. Anticancer agents are usually prepared as emulsions by manually mixing them with Lipiodol using a three-way stopcock without adding a surfactant. This yields a relatively unstable emulsion since a surfactant is required to stabilize the system. Body temperature may also facilitate aggregation of the emulsion, causing its breakdown in the blood stream after injection into the body. Once the emulsion breaks down, the anticancer agent and Lipiodol reach the tumor microcirculation as separate components, which does not allow Lipiodol to work as an efficient carrier. Miriplatin may be superior to other agents since it will be delivered to the tumor combined to Lipiodol. Moreover, compared to epirubicin or cisplatin, miriplatin induces lesser vascular damage.[11,12] TACE is indicated for BCLC intermediate stage, which is typically characterized by multiple lesions that require repeated treatment. It is reported that anticancer agents can cause arterial damage when injected into the hepatic artery. Additionally, they can cause occlusion of the vessel or arterioportal shunt. This compromises drug delivery following TACE sessions and may lead to insufficient drug accumulation in the tumor. Miriplatin may be thus suitable for repeated TACE.

The results of a previous randomized control trial showed that the antitumor effect of miriplatin is equal or superior to that of epirubicin or cisplatin. However, it is also reported that administering miriplatin as a high-viscosity formulation or large-droplet suspension may impede its delivery.[13–15] One solution to this problem is to use warmed miriplatin because the viscosity of miriplatin decreases on warming.[16] It has been demonstrated in some clinical studies that miriplatin shows a higher efficacy and improves tumor control better when it is warmed; however, local tumor control rate remains unsatisfactorily low (objective response rate: 40–46.7 %).[17,18] Moreover, the temperature of administered miriplatin suspension decreases along the way to the tumor, which attenuates the warming effect.

Another solution is to emulsify the miriplatin suspension. The droplet sizes of miriplatin suspensions vary, and occlusion of the feeding vessel can occur when large droplets are stacked in the proximal hepatic artery before optimal accumulation of the suspension is achieved.[15] This problem can be solved by using a monodisperse emulsion. Optimal accumulation of the emulsion can be obtained by carefully setting the droplet size of the emulsion to be less than the diameter of feeding arteries. The emulsion can also be stabilized by adding surfactant to the formulation. The results of the present study showed better Lipiodol accumulation in the tumors, both immediately after and two days after TACE, with the s/o/w emulsion than with the conventional suspension. Lipiodol retention in the tumors was also better with the s/o/w emulsion, which possibly reflects the stable nature of the emulsion. Moreover, necrosis of the tumor-normal parenchymal interface was more common in the experimental group. Local tumor recurrence tended to occur in the lesions and a sufficient margin is essential for tumor control when performing TACE.[19] Therefore, local tumor recurrence may be reduced by using the s/o/w emulsion.

No significant adverse events were observed in the experimental group; however, cholecystitis was observed in one rabbit in the control group. Lipiodol accumulation in the gallbladder wall was more common immediately after TACE in the experimental group; however, washout was observed in all rabbits in the group two days after the procedure. This may due to the stability and uniform droplet size of the emulsion. This is because if the emulsion is unstable, it can break down inside the arteries feeding normal structures. This will result in exposure of blood vessels to the anticancer drug, which may cause vascular injury. Ischemic injury may also occur if large Lipiodol droplets block proximal arteries; however, this is less likely to occur with the s/o/w emulsion than with the conventional suspension. Ischemic bile duct injury, which is a major complication and drawback of TACE, was not observed in this study. This indicates that the monodisperse s/o/w emulsion may be a safer alternative to the conventional suspension.

Promising results were obtained in the present study; however, a few clarifications are needed. Firstly, differences in the antitumor effects of the s/o/w emulsion and the conventional suspension are unclear. This is because the amounts of miriplatin and Lipiodol in the administered doses were lower than the required amounts for complete tumor necrosis. This was done to notice differences in Lipiodol accumulation in the tumors more clearly. Thus, the extent of necrosis should be evaluated in further studies after the full doses of the two ingredients are administered to both groups of rabbits. Additionally, the two-day period from TACE to animal sacrifice was too short for effective evaluation of antitumor effect. It has been shown that, in VX2-tumor-bearing rabbits, the difference in the concentration of an anticancer agent between the tumor and normal liver parenchyma peaks in approximately three days after TACE is performed when an emulsion prepared using SPG membrane is administered to the rabbits.[20] The main purpose of the present study was to evaluate the difference in Lipiodol distribution following treatment with conventional miriplatin suspension or miriplatin s/o/w emulsion. Thus, the observational period in future experiments should be longer. Furthermore, the droplet size of the emulsion should be considered. For instance, in the present study, an SPG membrane with a pore size of 20 μm was used to obtain an emulsion with a peak droplet diameter of 60 μm. This was because a previous clinical study on the efficacy of an epirubicin w/o/w emulsion revealed that local tumor control was better when droplet size 70 μm than when it was 30 μm.[21] Emulsions with small-sized droplets can enter microcirculation without being trapped, which indicates that droplet sizes as large as 70 μm are recommended. However, the data on the vessel diameters of rabbits are scarce and optimal target droplet size for rabbits should be determined by further investigation.

In conclusion, Lipiodol accumulation and retention in VX2 tumor may be improved by using a monodisperse emulsion, which is associated with no significant adverse events.

## Supporting information captions

**S1 Video. Injection of s/o/w emulsion.**

